# Dynamic readout of the Hh gradient in the Drosophila wing disc reveals pattern-specific tradeoffs between robustness and precision

**DOI:** 10.1101/2022.12.21.521489

**Authors:** Rosalío Reyes, Arthur D. Lander, Marcos Nahmad

**Affiliations:** Department of Physiology, Biophysics, and Neurosciences; Center for Research and Advanced Studies of the National Polytechnic Institute (Cinvestav-IPN); Mexico City, 07360; MÉXICO; Department of Developmental and Cell Biology; Center for Complex Biological Systems; University of California – Irvine, 92697; USA

## Abstract

How patterning by morphogen gradients determine tradeoffs between robustness and precision is unclear. Gradients that exhibit self-enhanced ligand degradation provide robustness to perturbations in morphogen production rates. However, increased robustness achieved through this mechanism is traded off for lower precision to noise. Here we use a hypotheses-driven theoretical approach to show that Hedge-hog (Hh) signaling would confer the same robustness to all target genes established by the steady-state gradient, but when a dynamical interpretation of patterning is used, robustness is maintained for steady-state outputs, but is traded off for higher precision in outputs set prior to steady-state. As predicted by our model, the widths of *decapentaplegic (dpp)*, and *collier (col)*, two target genes established by the Hh gradient in the *Drosophila* wing disc, exhibit differential robustness to Hh dosage. Particularly, higher robustness in the *col* pattern is ensured by Hh-dependent upregulation of its receptor Patched, an evolutionary-conserved property of Hh signaling that results in self-promoted Hh degradation. In contrast, *dpp* expression that is determined by the transient overshoot gradient, is insensitive to self-enhanced ligand degradation and exhibits less robustness, in exchange for a more precise boundary. Our work reveals of how morphogen gradients can establish tunable patterning properties in a target-specific manner.

## 1 Introduction

Developmental patterning needs to be robust to certain genetic and environmental perturbations in order to ensure a reproducible and functional body plan. Since patterns of gene expression are often specified by morphogen gradients, there has been considerable interest in understanding how these gradients robustly form and are interpreted (Neumann and Cohen, 1997; Gurdon and Bourillot, 2001; Lander, 2007; Claret et al., 2007; Ibanes and Belmonte, 2008; Rogers and Schier, 2011; Li et al., 2018; Stapornwongkul et al., 2018). The developing wing or wing imaginal disc of *Drosophila* has become a useful system to study the mechanisms of morphogen formation and interpretation (Hartl and Scott, 2014; Restrepo et al., 2014; Chen and Zou, 2019). Along the anterior-posterior (AP) axis, the *Drosophila* wing is patterned by morphogen gradients of Hedgehog (Hh) and Decapentaplegic (Dpp) that determine the position of the longitudinal veins L2-L5 (Blair, 2007). Hh is produced in cells of the posterior compartment of the wing imaginal disc and forms a short-range signaling gradient into the anterior compartment (Tabata and Kornberg, 1994). The Hh gradient organizes AP patterning of the wing both directly and indirectly; directly defines adult patterning outcomes, such as the expression of the transcription factor *knot* or *collier (col)* which sets the distance between the longitudinal veins L3 and L4 (Vervoort et al., 1999; Mohler et al., 2000); and also transcriptionally activates the expression of *decapentaplegic* (*dpp*) in a domain broader than *col* (Basler and Struhl, 1994; Vervoort, 2000). Dpp then acts as a long-range signal to globally coordinate patterning and growth along the AP axis of the wing disc (Affolter and Basler, 2007).

Contrary to other signaling pathways in which a ligand activates a signaling cascade by binding to its receptor, Hh signaling is activated by removing the receptor Patched (Ptc) from the plasma membrane, a process that is promoted by Hh binding and endocytosis (Torroja et al., 2005). This suggest that Hh signaling activity solely depends on the number of unbound Ptc receptors. However, a study suggested that the levels of Hh-bound Ptc can titrate the inhibitory effects of unbound Ptc and proposed that Hh signaling activity is more accurately represented by the ratio of bound to unbound Ptc receptor (Casali and Struhl, 2004). Importantly, a widely conserved feature of the Hh signaling pathway is that *ptc* is itself a target of the signal. Since Ptc expression attenuates the dispersion and strength of signaling activity, Hh-dependent Ptc upregulation acts as a negative feedback that plays important roles in self-limiting the range of the gradient (Chen and Struhl, 1996; Briscoe et al., 2001), desensitizing signal exposure over time (Dessaud et al., 2008), and controlling the spatial dynamics of the gradient (Nahmad and Stathopoulos, 2009). Additionally, this feedback property is a molecular implementation of self-enhanced ligand degradation which has been proposed to provide robustness of the Hh gradient to variations in Hh production rates (Eldar et al., 2003).

Hh signaling in the *Drosophila* wing disc exhibits two properties that determine its range and interpretation: (1) It limits its own range by promoting Hh ligand sequestration close to its source through the upregulation of its own receptor (Chen and Struhl, 1996); and, (2) it interprets positional information in a temporal manner using a single-threshold signaling range defined by a transient overshoot and the steady-state gradients (Nahmad and Stathopoulos, 2009). The first property is an experimental implementation of the self-enhanced ligand degradation strategy discussed above that exhibits robustness to changes in ligand production rates (Eldar et al., 2003; Lander et al., 2009). The second property determines the mechanism of gradient interpretation in which some patterning outputs (*dpp*) are established by an extended pre-steady state gradient, known as the *overshoot*, while others (*ptc* and *col*) are established by the steady-state gradient. Since the overshoot occurs prior to Hh-dependent Ptc up-regulation it benefit from the robustness offered by the self-enhanced ligand degradation mechanism.

A study by Irons *et al*., compared the width of *col* expression in the wing disc as well as the L3-L4 intervein distance in adult wings of *hh* heterozygous and wild-type animals and found that they are not statistically different, supporting that some robustness to Hh dosage is exhibited by the system (Irons et al., 2010). Recently, Hatori *et al*., showed that both width of *col* and *ptc* do not change in discs with 1, 2, 3 or 4 *hh* gene copies (Hatori et al., 2021). However, it remains unclear if the same robustness is exhibited by other Hh-dependent patterning outputs. Here we analyze morphogen diffusion models and show that if the patterns are established by the steady-state gradient all target genes are defined with the same robustness with respect to source variations, in agreement with prior theoretical work (Eldar et al., 2003). However, when the Hh gradient is interpreted dynamically through the overshoot model (Nahmad and Stathopoulos, 2009), robustness to *hh* dosage becomes target-specific. We show experimentally that the robustness of *col* pattern to changes in the number of *hh* copies is dependent on Hh-dependent Ptc upregulation. However, contrary to the steady-state interpretation, the *dpp* pattern is much less robust to changes in Hh levels, *i*.*e*., the robustness to Hh levels in this system is target specific. We show that this reduced robustness in *dpp* patterning is compensated by an increased precision of the *dpp* border. Therefore, our work shows that patterning system driven by a single morphogen can be tuned to exhibit different robustness and precision properties.

## 2 Results

### 2.1 Steady-state interpretation of morphogen gradients reveals the same robustness to morphogen dosage for all targets

Prior work on morphogen robustness has relied on quantifying changes to either on overall gradient shape (Gurdon and Bourillot, 2001; Tabata and Takei, 2004) or to a single threshold location defined by a specific length-scale of the gradient (Eldar et al., 2003). The underlying assumption is that robustness is the same for all targets, but this has not been studied in detail. To address this, we investigated robustness predicted by classical gradient models, where the steady-state morphogen gradient defines territories at specific thresholds. As a starting model, we considered the steady-state, free-diffusion, linear-degradation model in which the source of the morphogen is a boundary condition (see Box 1 and Supplementary Information). In this case, perturbations in the morphogen source induce a uniform displacement of the profile and the territories defined by the morphogen, (figure 1A-C), suggesting that different patterns exhibit the same response to perturbations. This is consistent with previous work by Eldar *et al*., which shows that a more general, non-linear degradation model at the steady-state also displaces all territories by the same amount upon perturbations in the morphogen source (Eldar et al. (2003); Box 1 and Supplementary Information). This occurs because the solution to the perturbed problem is a displacement of the profile with a new boundary condition (see Box 1).

**Figure 1:**
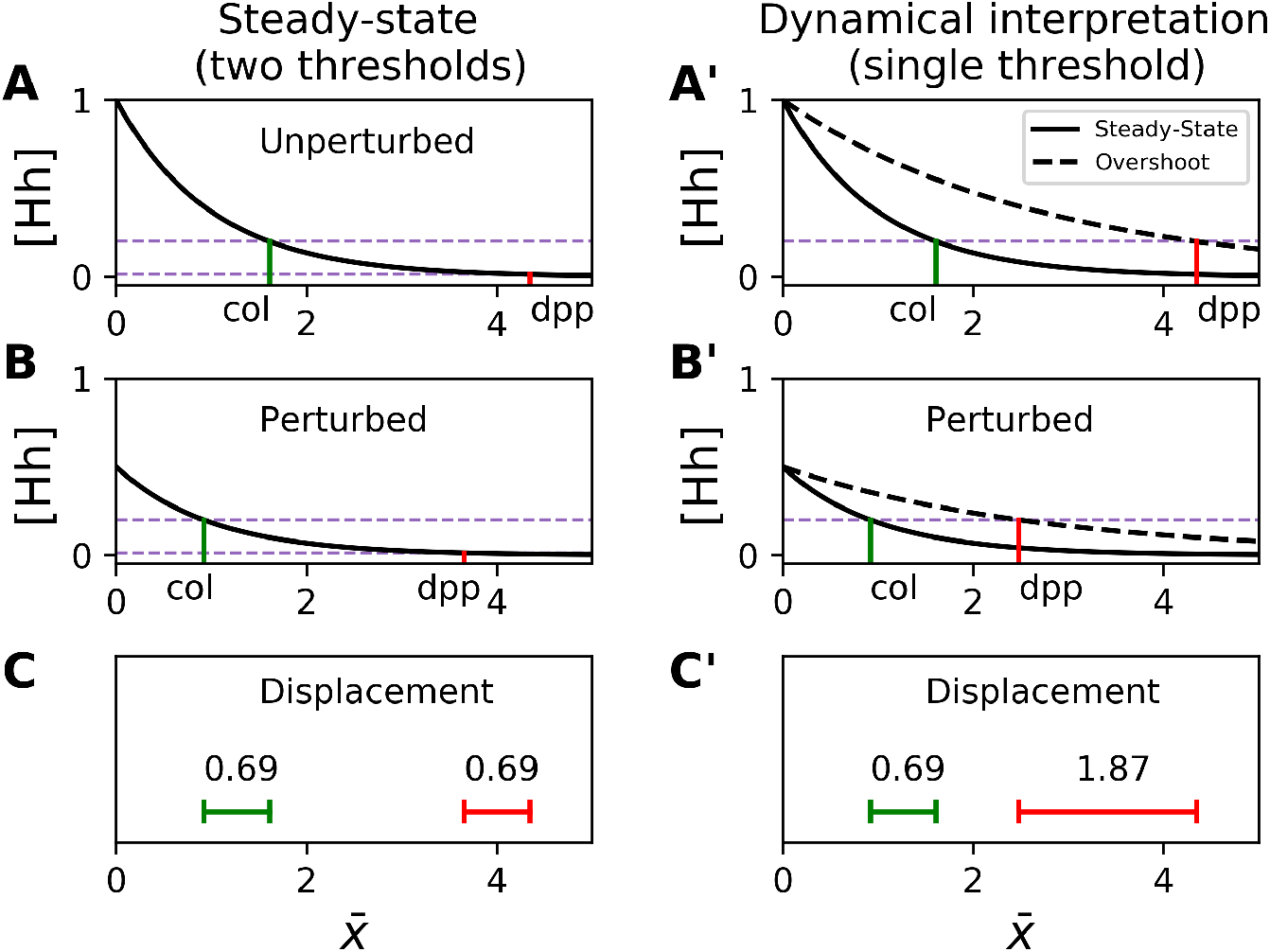
Dynamical interpretation model reveals differential robustness to reducing morphogen dosage to half. Numerical solution of equation (6) (Box 2) using full (*Hh*(*x* = 0) = 1; A,A’) or half (*Hh*(*x* = 0) = 0.5; B, b’) dosage of Hh. The positional shift for the classical (steady-state; A-B) model with two threshold (corresponding to *col* and *dpp*) is identical (C); whereas the dynamical interpretation model with a single threshold readout using the overshoot vs. the steady-state gradient predicts different shifts (C’). We consider for this graphs *β*_*late*_ = 1, *β*_*early*_ = 1*/*9 and *D* = 1, that implies *λ*_*SS*_ = 1 and *λ*_*pseudoSS*_ = 3*λ*_*realSS*_. 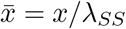.

Next, we considered a model in which the source is not modeled as a boundary condition, but instead, as a domain with constant production of the ligand. Indeed, a simple-diffusion model with a source can be reduced to a model with the source as a boundary condition (Box 1 and Supplementary Information).

We then expanded this defined-source model to a case that approximates the Hh morphogen system in the *Drosophila* wing. Since Ptc is upregulated by Hh signaling and contributes to Hh degradation by binding the Hh lig- and, we considered one-dimensional model in which ligand degradation has different values within and beyond the Ptc expression domain (see Box 1). Consistent with the results of the previous models, any two target genes whose borders are defined by different concentration thresholds will exhibit the same robustness (Box 1 and Supplementary Information). Taken together, we conclude that classical models of morphogen interpretation at the steady-state exhibit the same robustness of multiple target genes defined by Hh.

#### Box 1.

Steady-State models reveals the same robustness for all target genes

The simplest model of a morphogen is the one that considers simple diffusion and linear degradation. The steady-state model equation is *D*∇^2^*M* − *βM* = 0, where *λ*^2^ = *D/β* is the ratio between the rate of ligand degradation *β*, and the diffusion coefficient D (Eldar et al., 2003). Upon source perturbations 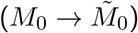 all pattern boundaries defined at different thresholds will move uniformly by the amount:

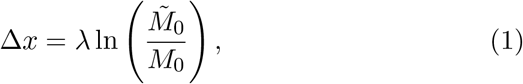

The result holds even if morphogen degradation is considered an arbitrary function depending only on *M*, the concentration of the ligand (Eldar et al. (2003) and Supplementary Information).

##### Basic Diffusion-Degradation Model with a non-point source of lig- and production

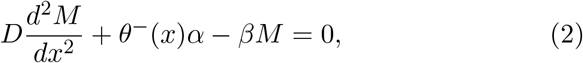

where *θ*^−^(*x*) = 1 for *x* < 0 and zero otherwise, *α* is the rate of ligand production and *β* the rate of ligand degradation. If we assume the function is continuous at the boundary (*x* = 0), the solution reduces to:

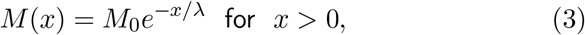

where 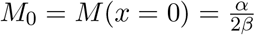. Then any non-point source can be reduced to a point source, and equation (1) holds when the morphogen is not a boundary condition.

##### Simplified model of Hh signaling

We also considered a simple model of the mode Hh gradient, where Hh is produced in the negative compartment (*x* < 0) and is degraded at two different rates in the positive compartment (*x >* 0):

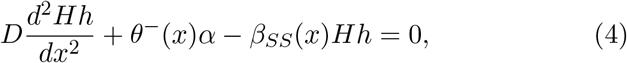

where *β*_*SS*_(*x*) ≈ *β*^*^ for 0 < *x* < *b* (*b* is the width of Ptc stripe) and *β*_*SS*_(*x*) ≈ *β* for *x* ≥ *b*. The solution is given by:

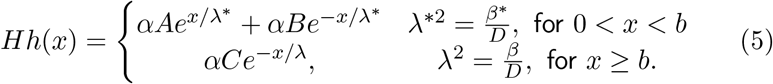

where *A, B*, and *C* are parameters defined by imposing continuity of the function at *x* = 0 and *x* = *b* (see Supplementary Information for additional details).

### 2.2 Mathematical analysis shows that a dynamical interpretation of Hh signaling implies differential robustness

Previous work suggest that Hh signaling is interpreted dynamically in the *Drosophila* wing disc (Nahmad and Stathopoulos, 2009). This dynamical model proposes that because of Hh-dependent Ptc upregulation, the Hh gradient has a maximal reach (which is wider than the steady-state), thus defining two characteristic signaling times with a single threshold: the transient overshoot and the steady-state gradient (Nahmad and Stathopoulos, 2009). We then analyzed a simplified dynamic model of Hh diffusion (see Box 2 and Supplementary Information) that takes into account the temporal upregulation of *ptc* through a temporal degradation step-wise function. The model shows that if the time of change of degradation is long enough, there will be transient solutions with a greater reach than the steady-state. Following Nahmad and Stathopoulos 2009 we define the overshoot as the transient gradient of maximum reach (figure 1B’). A robustness analysis shows that territories defined by the overshoot are less robust than territories defined by the steady-state (see Box 2, figure 1B’ and Supplementary Information). Then, in contrast to the classical model, the dynamical interpretation model predicts differences in target gene displacement upon perturbation in morphogen dosages (figure 1C,C’). In particular, the pattern established by the overshoot does not exhibit the robustness of the pattern defined by the steady-state (figure 1C’), as expected, because it is established before the effect of self-enhanced ligand degradation (*β*_*early*_ < *β*_*late*_; see Box 2).

#### Box 2.

Dynamic models implies differential robustness

##### Simplified dynamic model of Hh signaling

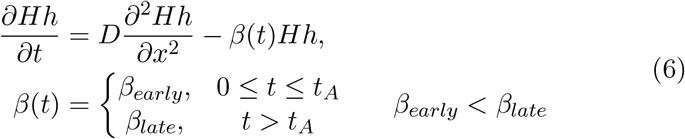

where *t*_*A*_ is the time at which the rate of degradation changes between initial rate *β*_*ealy*_ and final rate *β*_*late*_. Then, overshoot is delimited by two steady-state (SS) solutions, the pseudo-SS solution correspondent to *β*_*early*_,

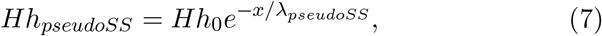

and the real steady-state solution (real-SS) correspondent to *β*_*late*_,

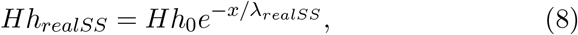

where *Hh*_0_ is the concentration of morphogen at *x* = 0. Since *β*_*early*_ < *β*_*late*_ (6), then:

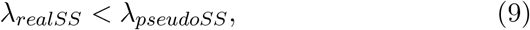

Hence, *Hh*_*pseudoSS*_ has a greater reach than *Hh*_*realSS*_.

We define the *coefficient of robustness* as

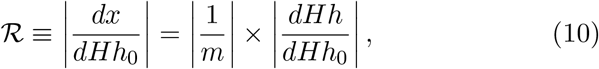

where *m* is the slope of Hh at the border of the defined territory. The smaller the robustness coefficient, the more robust the system is. Since for our model (6), 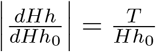, where *T* is the threshold concentration that defines the territory, then, ℛ ∝ 1*/*|*m*|. From equations (7) and (8) we obtain

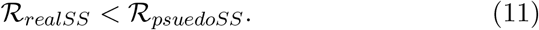

Taken in to account continuity of solutions we conclude that ℛ_*realSS*_ < ℛ_*overshoot*_ < ℛ_*psuedoSS*_ (see Supplementary Information), *e. i*., the overshoot is less robust than steady-state.

### 2.3 The target defined by the overshoot trades off robustness for precision

The finding that target genes defined by the overshoot are less robust than in the steady-state model could be seen as a disadvantage of the dynamic interpretation model, since one might expect that biological processes always seek greater robustness. However, robustness is just one patterning property. Morphogen concentrations are naturally noisy, which causes territories to be defined by a diffuse border (Lander et al., 2009). A previous study has shown that noise is attenuated at steeper morphogen slopes, but is amplified as the morphogen flattens (Gregor et al. (2007), figure 2A). Therefore, precision is directly related to the slope of the morphogen gradient at the position where the pattern is established. If we define 𝒫, the *precision coefficient* (Box 2), as the absolute value of the rate of change of position *x* upon fluctuations on gradient concentration (which is the same that the absolute value of the inverse of the gradient slope), 𝒫 ≈ 0 corresponds to locations of low precision (sharp slope). Hh-dependent Ptc upregulation causes a retraction of the gradient and a steep slope (𝒫 ≈ 0), defining a sharp boundary of gene expression. However, for targets specified several cells away from the morphogen source, the steady-state gradient has a much flatter slope than the slope of the overshoot gradient at that location (figure 2B, B’), indicating that 𝒫 _*steady*−*state*_ *>* 𝒫 _*overshoot*_, *i*.*e*., beyond a certain location (see Supplementary Information and Supplementary figure 1), the dynamic interpretation is less robust to ligand dosage, but offers more precision to noise than the steady-state gradient at lower threshold (figure 2B,B’)

**Figure 2:**
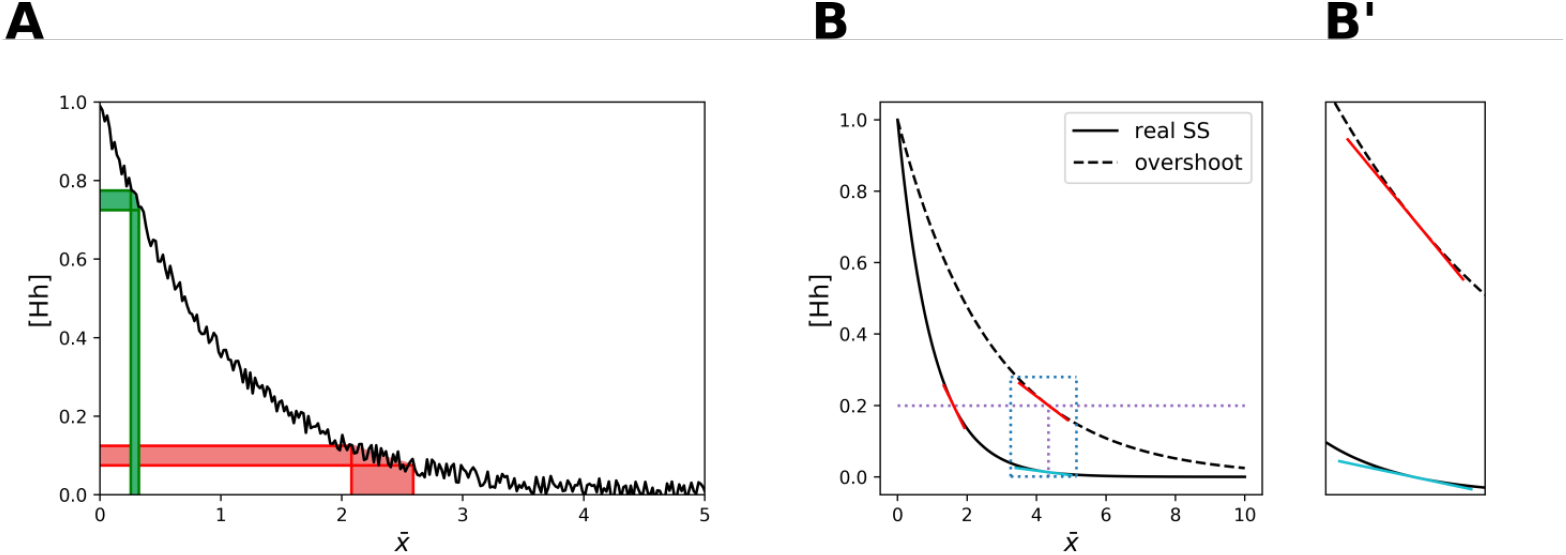
Precision to noise drops considerably with distance, but is much improved if patterning is established by the overshoot gradient. A. Accuracy of positional specification under uniform noise of amplitude 0.05 using two random thresholds in the steady-state solution of the model in Box 2. B. Comparison of slopes (that define the precision coefficient 𝒫) during the dynamical interpretation (red lines) using a single threshold of 0.2 vs. the corresponding slope that would define that position with the steady-state gradient (cyan line). B’. Inset of the blue dotted rectangle in B comparing the precision coefficient of the overshoot vs. steady-state gradient at the position defined by the overshoot.

### 2.4 Simulations in a more explicit model of Hh signaling exhibit Hh-dosage robustness to patterns established by the steady-state, but fragility of patterns established by the overshoot

We then asked if our results of the previous sections also hold in a dynamic model of the Hh pathway presented in Nahmad and Stathopoulos, 2009; (see equations (12)-(16), Box 3). We solved the equations numerically and computed the expected shift in the establishment of the overshoot and steady-state targets upon perturbations of the Hh production rate, *α*_*Hh*_, in the posterior compartment. Indeed, if we gradually upregulate or down-regulate *α*_*Hh*_, the steady-state outputs are more robust than the overshoot outputs of the signal, (figure 3A). The same result is obtained when we define the *Coefficient of Displacement* (CD) as the average positional displacement (at the concentration threshold of 0.2), resulting from varying *α*_*Hh*_ by half or twice of its wild-type value (see equations (18) and (19) in Materials and Methods), which allows 9 comparisons with experimental manipulations. Moreover, the excess in CD defined by the overshoot relative to the steady-state gradient do not depend on the specific choice of parameters in the model (figure 3B). We conclude that under the interpretation of overshoot model, our model predicts higher robustness for targets specified by the steady-state gradient (*col*), with respect to those specified by the overshoot profile (*dpp*).

**Figure 3:**
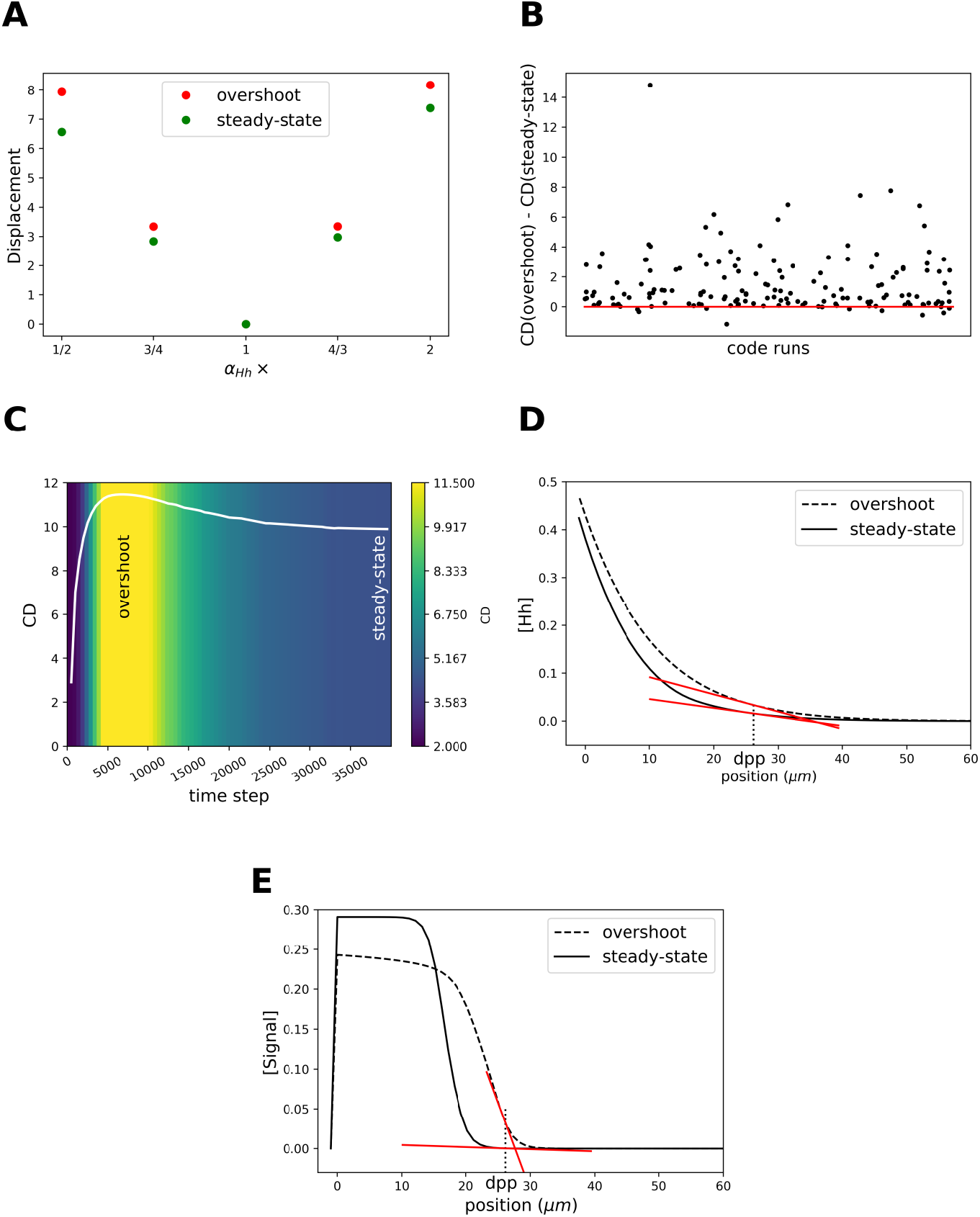
Analysis of robustness and precision in a more explicit model of Hh signaling confirms our prior findings. A. Displacement, δ_*q*_ (see Material and Methods) of the positional specification for the overshoot (red) vs. steady-state (green) signal outputs upon gradual variations in *α*_*Hh*_ using the original parameters defined in Nahmad and Stathopoulos, 2009. B. Differences in the Coefficient of Displacement (CD; see Materials and Methods) between the overshoot and the steady-state gradients when all parameters in the system are varied between 0.5 and 2.0 of the reported values in Nahmad and Stathopoulos 2009. C. CD of Hh signaling as a function of time. D, E. Precision coefficients obtained from the tangent lines to the Hh profiles (D) at the overshoot and steady-state and Signal (E).

Prior theoretical work showing that the interpretation of positional information prior to steady-state actually improves robustness (Bergmann et al., 2007), appears to contradicts our finding that overshoot dependent targets (which occurs prior to steady-state) are less robust than steady-state-dependent targets (figure 3A,B). In order to understand the relative robustness of pre-steady state gradients in the dynamical model considered here, we computed the CD as a function of time in our model of Hh signaling (see Supplementary Information). We found that early transient states exhibit the smallest CDs and therefore are the most robust among all the states (figure 3C), in agreement with prior studies (Bergmann et al., 2007), but the CD increases as the gradient approaches the overshoot when the CD reaches a maximum, before it starts to decrease again towards the steady-state (figure 3C).

In addition, we wanted to confirmed the result of the previous section in the whole Hh signaling model, so we analyzed the precision coefficient, 𝒫, for both the morphogen degradation profile, *Hh*(*x*), and the activation signal profile, *Signal*(*x*). We find that at the approximate position where *dpp* is defined, 𝒫 of the Hh steady-state is approximately twice as 𝒫 of the Hh overshoot (figure 3D). Surprisingly for Signal, the steady-state profile is 82 times more imprecise than the Signal overshoot at the *dpp* border (figure 3E). Taken together, we conclude that the overshoot model recapitulates the differential robustness of Hh target genes observed in the simplified dynamical model (figure 1A’-C’) while allowing the *dpp* edge to be defined more precisely than if it were defined by the steady-state.

#### Box 3.

Hh signaling model

A more explicit model of the Hh pathway that includes receptor binding, upregulation and signaling can be described by the following set of reaction-diffusion equations (Nahmad and Stathopoulos, 2009)

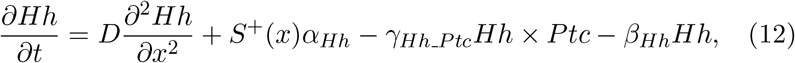

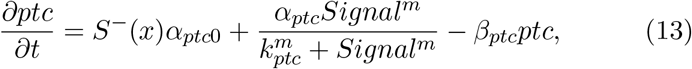

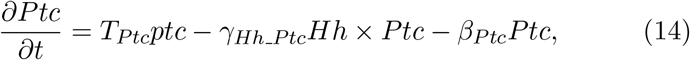

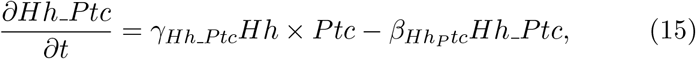

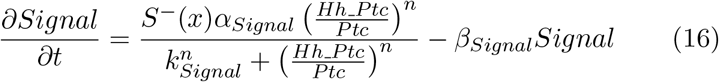

where [*Hh*], [*ptc*], [*Ptc*], and [*Hh Ptc*] are the concentrations of Hh, *ptc* (mRNA), Ptc (protein) and the Hh-Ptc complex, respectively. The coefficients *α, β, γ*, and *T* represent the rates of synthesis, degradation, complex formation, and translation, respectively. We used a system of coordinates centered on the AP boundary with the anterior compartment on the negative side (left). *S*^+^(*x*) (or*S*^−^(*x*)) is a step function of the form *S*^+^(*x*) = 1 if *x >* 0 [or *S*^−^(*x*) = 1 if *x* < 0] and zero otherwise. [*Signal*] represents the concentration of Hh signaling activity and assume that [*Signal*] reflects the intracellular response to activate Hh target gene expression. The system (12-16) is subject to the following boundary and initial conditions:

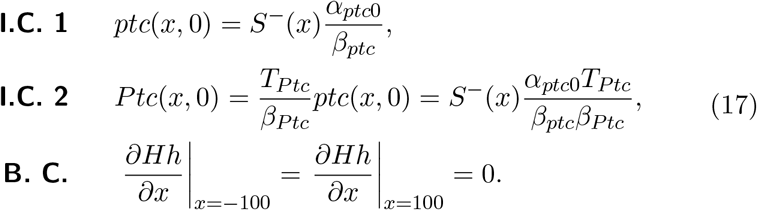

### 2.5 Robustness of steady-state outputs depends on Hh-dependent *ptc* regulation

Previous work suggests that Hh-dependent *ptc* upregulation self-regulates the range of the signal (Chen and Struhl, 1996). We asked to what extent Hh-dependent *ptc* regulation contributes to the robustness of patterning outputs. We gradually modulated the *ptc* production rate, *α*_*ptc*_, and noticed that the CD of the steady-state is greatly reduced with *α*_*ptc*_, but has little effect on the CD of the overshoot, (figure 4A). Once again, this result is independent of the choice of parameters, (figure 4B). We conclude that Hh-dependent *ptc*-regulation provides robustness to steady-state but has little effect in the overshoot output. Therefore, we suggest that Hh-dependent Ptc upregulation explains the differential robustness in this system by making steady-state outputs more robust while leaving overshoot-defined outputs unchanged.

**Figure 4:**
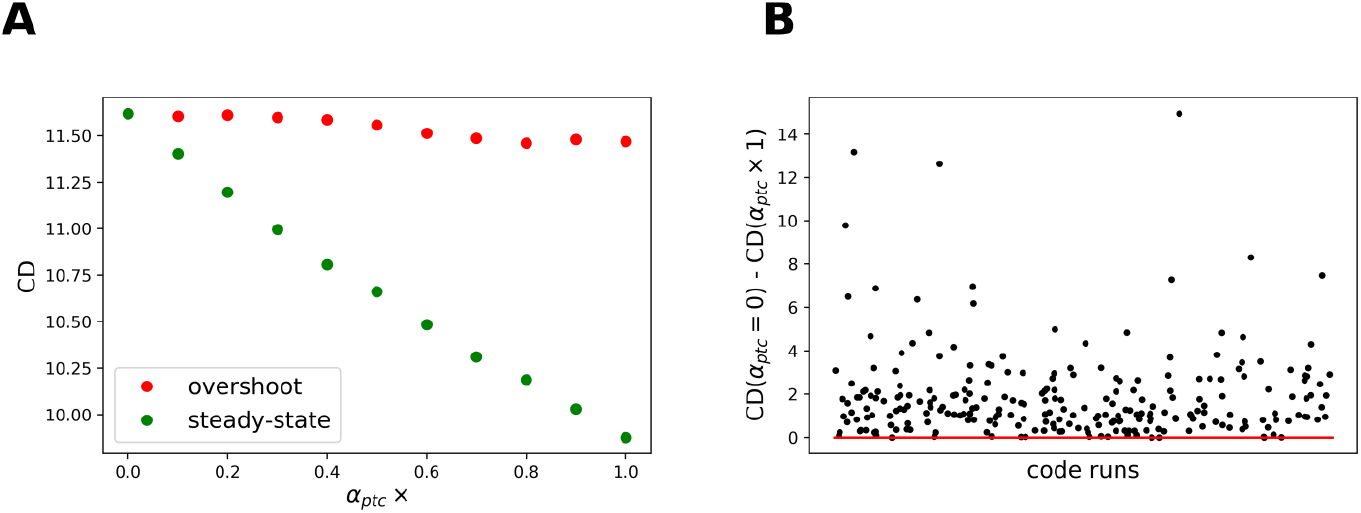
Simulations reveal that steady-state robustness depends on *ptc*-regulation. A. Computation of CD upon reduction of *α*_*ptc*_ relative to the reported value for steady-state and overshoot outputs. B. Dependence of steady-state robustness (as defined by the difference in CD with (*α*_*ptc*_) and without (*α*_*ptc*_ = 0) on Hh-dependent *ptc* upregulation when the rest of the parameters are varied.

### 2.6 Unlike *col*, the pattern of *dpp* does not exhibit robustness to changes in Hh levels

We then proceeded to test experimentally whether Hh targets are differentially robust to changes in Hh dosage as predicted by the dynamical interpretation model. Previous study showed that the width of the *col* domain is unaffected in *hh* heterozygous wing discs (Irons et al., 2010; Hatori et al., 2021). To determine if this robustness property holds for *dpp*, which is established by the overshoot (Nahmad and Stathopoulos, 2009), we examined the patterns of *col* and *dpp* (using a lacZ reporter) in discs carrying 1 or 2 copies of *hh*. We show the width of *col* pattern in the heterozygote mutant disc is reduced by 1.64 *μm* (less than the width of a cell, 2.5 *μm*) relative to the width of the wildtype pattern, (figure 5A-B). While the heterozygote pattern of *dpp* is reduced by 5.56 *μm* (the width of two cells) relative to wildtype, (figure 5C-D). All these results are in agreement with our theoretical results from sections 2.2 and 2.4. Taken together we conclude that *col* pattern is more robust than *dpp* pattern.

**Figure 5:**
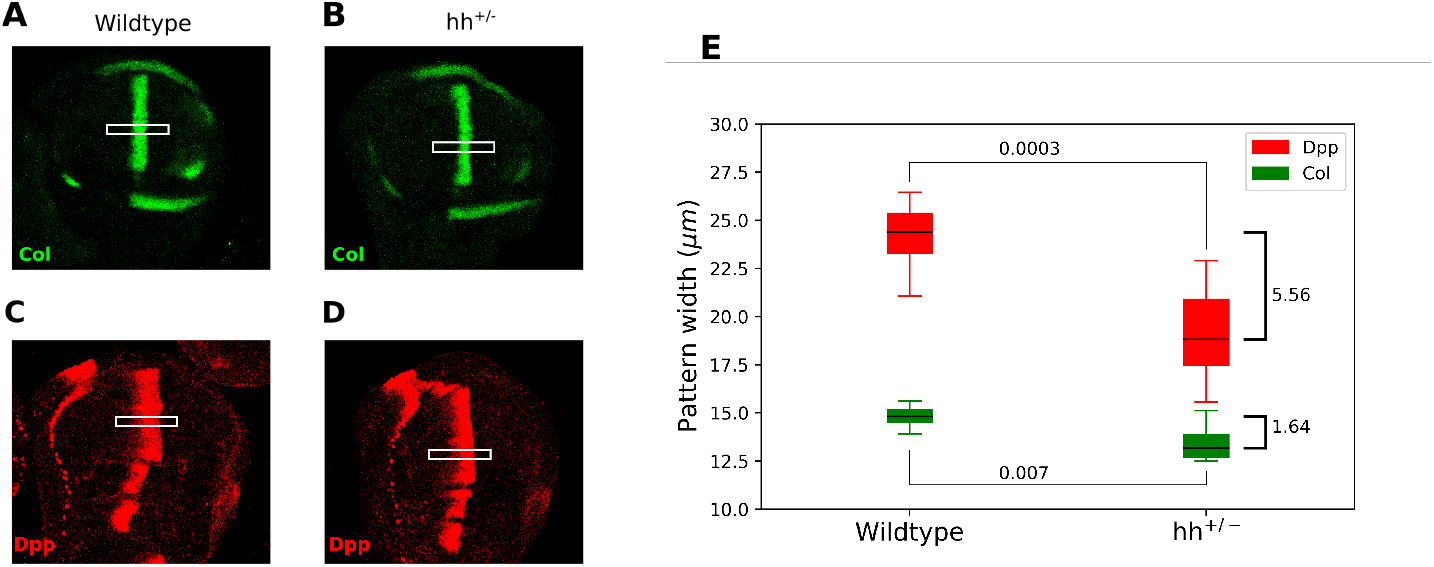
Differential robustness of Hh targets to *hh* dosage. Third-instar wild-type (B, D) and *hh* heterozygous (*hh*+/-) wing discs immunostained with Col (A, B) and *β*-galactosidase (C, D) antibodies. Both wild-type and *hh*+/- flies carry a transgene with a dppLacZ enhancer trap, so *β*- galactosidase marks the pattern of *dpp* expression. (E) Width of the gene pattern measured in the marked white rectangle. Marks on the right represent the difference between the medians of both groups.

### 2.7 Robustness of *col* expression depends on Hh-dependent Ptc upregulation

We then asked whether the robustness of the Col domain can be explained by receptor-enhanced ligand degradation as proposed in our theoretical studies and another works (Eldar et al., 2003; Irons et al., 2010). We considered *ptc* mutant flies that contain a transgene that expresses *ptc* under a constitutive promoter (Tubulin1*α >ptc>*Tubulin3’UTR or TPT; Chen and Struhl (1996)). *ptc*-/-; TPT/+ flies do not express Hh signaling-dependent *ptc*, yet these flies are viable, fertile, and apparently normal. Nonetheless, the expression patterns of *col* and *dpp* nearly overlap in *ptc*-/-; TPT/+ discs (Nahmad and Stathopoulos, 2009). If the robustness of *col* to Hh levels relies on the feedback between Hh signaling and Ptc, then the robustness should be broken in a *ptc*-/-; TPT background. We aimed to compare the patterns of expression in *ptc*-/-; TPT/*hh*AC larvae do not grow and die during the first instar (data not shown). However, *ptc*-/-; TPT/*hh*^*ts*2^ larvae, where *hh*^*ts*2^ is a sensitive allele which reduce the functionality of *hh* at temperatures higher than 25 °C but is unaffected at 18 °C (Ma et al., 1993), do survive and reach adulthood. Then, we analyzed *ptc*-/-; TPT/*hh*^*ts*2^ discs at 18 °C and 25-29 °C. We notice that in *ptc*-/-; TPT/*hh*^*ts*2^ discs, difference in medians at 18 °*C* and 25-29 °*C* is 4.57 *μm*, a value comparable to *dpp* pattern change under *hh* dosage changes (figure 5E). This is not due to an effect of temperature since the width of the Hh-dependent patterns appear to be insensitive to temperature (Supplementary figure 2). This result is in agreement with our theoretical result from section 2.5. Taken together, our work suggests that signal-dependent Ptc upregulation is responsible for providing robustness of Col expression to Hh dosage.

## 3 Discussion

The robust architecture of body plans to genetic and environmental perturbations is a general feature of developmental systems (Waddington, 1942; Csete and Doyle, 2002; Kitano, 2004). At the same time, this robust design should also admit some flexibility in order to allow the system to evolve into new phenotypes under certain genetic or environmental challenges (Barkai and Shilo, 2007). While much work has devoted to the understanding of which network features confer robustness in developmental patterning, it is unclear how a robust, yet flexible architecture could be encoded in the interpretation of morphogen gradients (Lander et al., 2009). In particular, the existence of differential robustness, *i*.*e*., the ability of a single morphogen to generate outputs that are more robust than others under certain perturbations has not been documented.

Relative to the classical view of morphogen interpretation, in which different concentration thresholds at the steady-state define different borders of gene expression patterns, two strategies have been proposed to increase robustness to changes in the rates of morphogen production. First, morphogen gradients that promote their own degradation and decay sharply near the source of ligand production (Eldar et al., 2003). And second, gradients that specify patterns prior to steady-state (Bergmann et al., 2007). When implementing either of these strategies, the same robustness is expected for all target gene expression, regardless of the concentration thresholds at which it is established (see Box 1). Both of these strategies trade off robustness at the expense of narrower gradients with respect to the classical steady-state gradient interpretation, and both not seem to provide a robust-yet-flexible design that could trade off some robustness in exchange for any another advantageous properties.

As we show here, experimental examination of *col* and a *dpp* reporter under different Hh dosage condition (*hh×*1 vs. *hh×*2) shows that *col* expression is more robust to changes in Hh dosage than the expression of the *dpp* reporter. Furthermore, the unique robustness of the *col* pattern is indeed enabled by Hh-dependent *ptc*-upregulation, a form of self-enhanced ligand degradation. The differential behaviour of *col* and *dpp* is inconsistent with a steady-state model, but is predicted by analytical and numerical exploration of a mathematical model in which *dpp* responds to the overshoot, and not the steady-state Hh gradient, a result that provides further support to the overshoot model (Nahmad and Stathopoulos, 2009).

Why does the system use this differential robustness patterning strategy? What would be the advantage of having a robust *col* pattern vs. a less robust *dpp* pattern? The *col* expression pattern defines a specific feature in the adult wing, the L3-L4 intervein area, which corresponds to the more central area of the wing, whereas the *dpp* pattern simply works as the source of another morphogen. As suggested by prior theoretical work, the source width of a morphogen does not have a significant impact on patterning (Mizutani et al., 2006), so perhaps the robustness of the *dpp* pattern was not subject to strong selection pressure during evolution, or there are other mechanisms that provide robustness at the level of Dpp signaling (Aguilar-Hidalgo et al., 2018; Romanova-Michaelides et al., 2022). On the other hand, this patterning strategy provides an additional advantage. While having *dpp* respond to the steady-state, rather than the overshoot behavior of the Hh gradient, could have conferred on *dpp* the same robustness to Hh dosage as Col, it would have come at a cost. This is because self-enhanced ligand-degradation gradients dramatically lose precision several lengthscales away from the source. By interpreting the *dpp* pattern using the overshoot of the Hh gradient, loss of robustness is offset by gains in precision, allowing the *dpp* stripe to be sharper than would have been the case otherwise (see Box 2 and figure 7). Taken together, our work provides further support to a dynamic interpretation of morphogen gradients and exemplifies how morphogen gradients may be wired to support patterns that need to be robustly established, while leaving others more flexible but with higher precision.

## 4 Materials and Methods

### 4.1 Fly stocks and crosses

Fly crosses were conducted at 25°C, except where otherwise indicated. For experiments using the *hh*+/- allele (figure 5A,C) fly stock *ry*[506]*hh*[AC]/TM3,Sb[1] (BDSC #1749) was crossed to fly stock *dpp*10638/CyO at 25 °C to obtain *dpp*10638; *ry*[506]*hh*[AC] discs. *hh*[AC] is a lost of function hh allele and *dpp*10638 is a transgene on II containing a lacZ reporter that produces nuclear -gal. A Tubulin1*>ptc>*Tubulin1 39’UTR (TPT) transgenic line located on chromosome III has previously been shown to rescue *ptc* mutant animals (Chen and Struhl, 1996). To obtain *ptc*-/-; TPT/*hh*^*ts*2^ (figure 6A,B) discs we crossed *ptc*^9^; TPT/SM6; TM6B, Tb males to *ptc*^16^/SM6; *hh*^*ts*2^/TM6B, Tb females. *ptc*^9^ produces a product that is defective in reaching the plasma membrane and binding to Hh (Strutt et al., 2001). *ptc*^16^ has been previously characterized as a null allele (Capdevila et al., 1994) and *hh*^*ts*2^ is a sensitive allele which reduce the functionality of *hh* at temperatures higher than 25 °C but is unaffected at 18 °C (Ma et al., 1993). To obtain *dpp*Gal4,UAS-GFP discs (Supplementary figure 2) *dpp*Gal4, UAS-GFP/[TM6B, Tb] (BDSC #84316) male was crossed to WT female.

**Figure 6:**
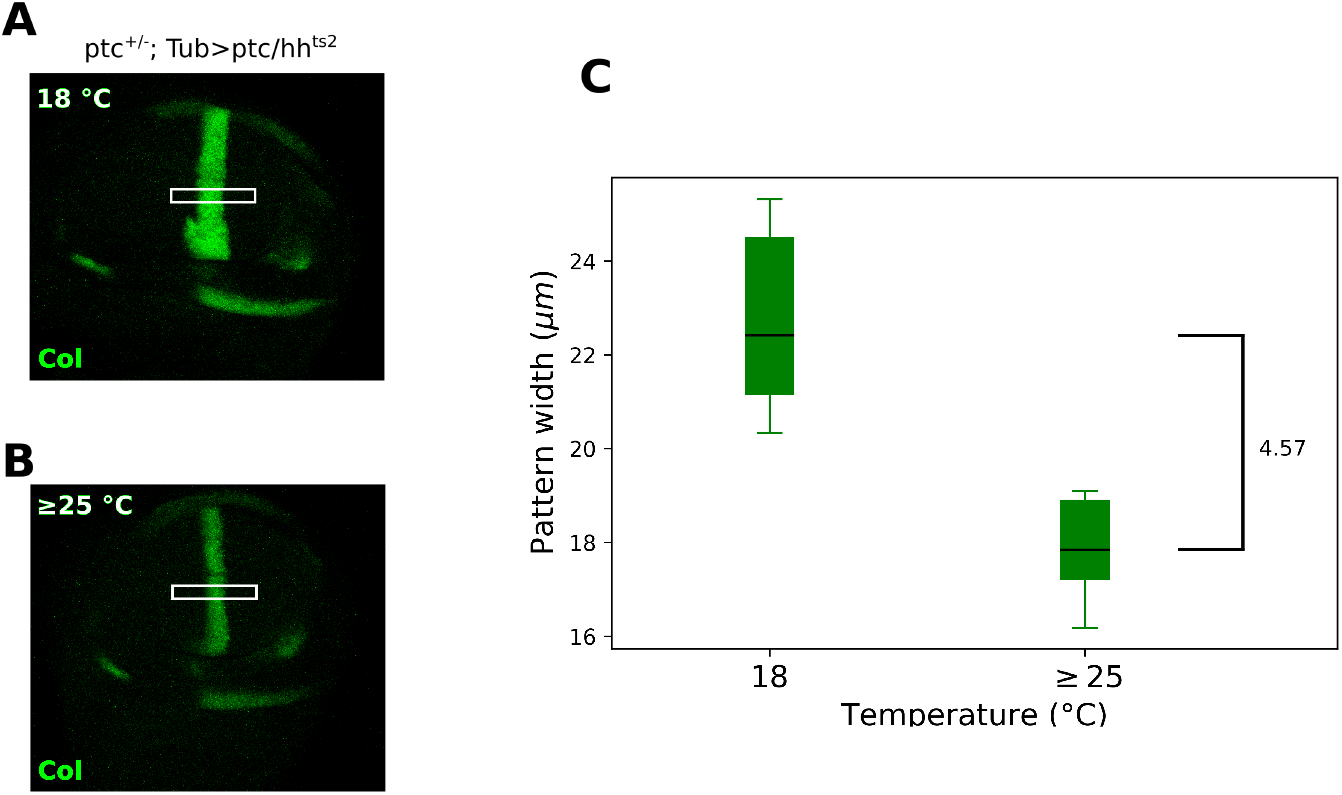
Col robustness depends on Ptc regulation. (A) *col* pattern in wing disc at 18 °C (both alleles are functional); (B) *col* pattern at 25 and 29 °C (only one allele is functional). (C) Width patterns measures in the market white rectangle. Mark on the right represent the difference between the medians of both groups.

**Figure 7:**
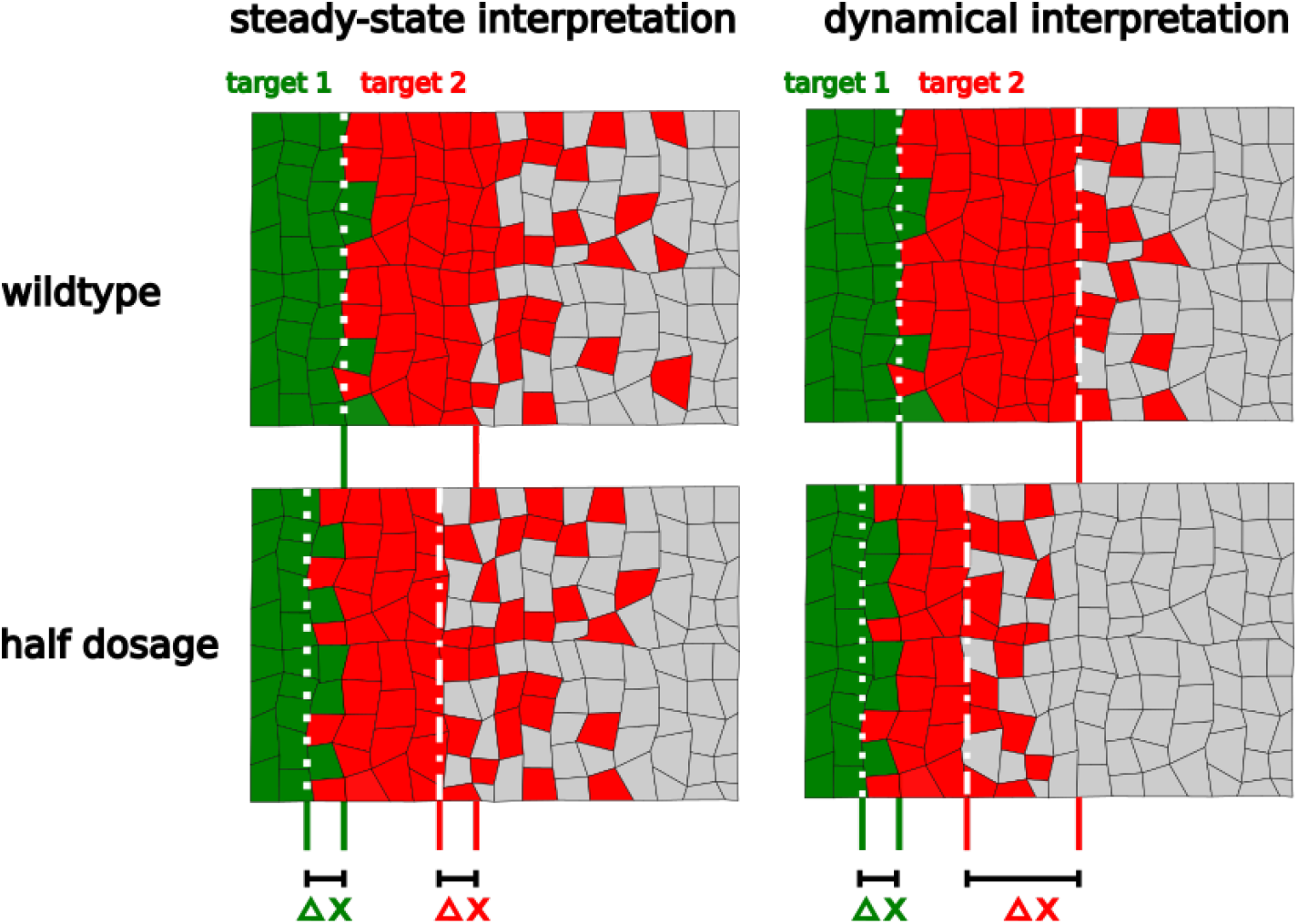
Scheme summarizing the results of this study. In the steady-state interpretation, all the target genes are established with the same robustness (Δ*x*) upon perturbations in the amount of ligand. In the dynamic interpretation one of the target genes (red) is defined with less robustness than the other (green). However, it allows the less robust gene to be defined with greater precision than the steady-state would define it (compare the widths of the transition zone, red-cell-grey-cell).

### 4.2 Wing imaginal disc dissection and immunostaining

Wing imaginal discs were dissected from third-instar larvae of both sexes. After dissection in a stereoscopic microscope, discs were fixed in PEM-T (PEM with 0.1% of Triton X-100) with 4% paraformaldehyde, washed 3 times and blocked in PEM-T with 0.5% of BSA (Bovine Serum Albumin) for 2 h at room temperature. Then, samples were stained with primary antibodies at 4 °C overnight at the following dilutions: monoclonal mouse anti-Col (M. Crozatier, 1:250), rabbit anti–gal (Invitrogen Molecular Probes, 1:250), monoclonal mouse anti-Ptc (developed by I. Guerrero, and was obtained from the Developmental Studies Hybridoma Bank at the University of Iowa, 1:250). Primary antibodies were detected with Alexa Fluor 488 anti-mouse and Alexa Fluor 555 antirabbit and anti-mouse Alexa Fluor 647 (1:1000). Imaging was done with a confocal microscope using a 40X oil-immersion objective.

### 4.3 Displacement and robustness measurements

In order to measure the displacement upon a perturbation in *α*_*Hh*_ we define

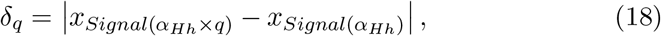

where *x*_*Signal*_ means the reach of Signal at 0.2 of maximum concentration and *q* means the factor by which the *α*_*Hh*_ was modified from its original value. In addition, we define the Coefficient of Displacement (CD) as:

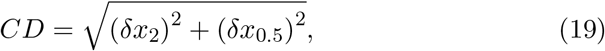

namely, an average measure of the change in signal reach when we change *α*_*Hh*_ by two fold of its ‘wild-type’ value (as reported in Nahmad and Stathopoulos, 2009).

### 4.4 Numerical simulations

A Forward-in-Time-Centered-in-Space (FTCS) algorithm (*x* = 1*μm, t* = 0.5*s*) was implemented to solve Supplementary Equations (25)-(29) in Python using the parameters reported by Nahmad and Stathopoulos 2009. The steady-state solution to Supplementary Equation (33) was solved using solve bvp from scipy.integrate Python package. Plots were made with matplotlib library of Python.

### 4.5 Image analysis

We take the Z projection of the images using ImageJ. We reduced background noise by subtracting the average intensity in a region outside the region of interest using ImageJ. The analyzed images were saved in png format, to then measure the width of the patterns using OpenCv library of python. We normalized the intensity values after dividing them by the maximum intensity, then we measured the width of a pattern stripe at 0.2 of relative intensity. These steps were coded in Python. Graphs were made with matplotlib and seaborn libraries of Python.

## 5 Acknowledgements

We thank M. Crozatier for kindly providing us with an aliquot of Collier antibody

## 7 Supplementary figures

**Supplementary figure 1:**
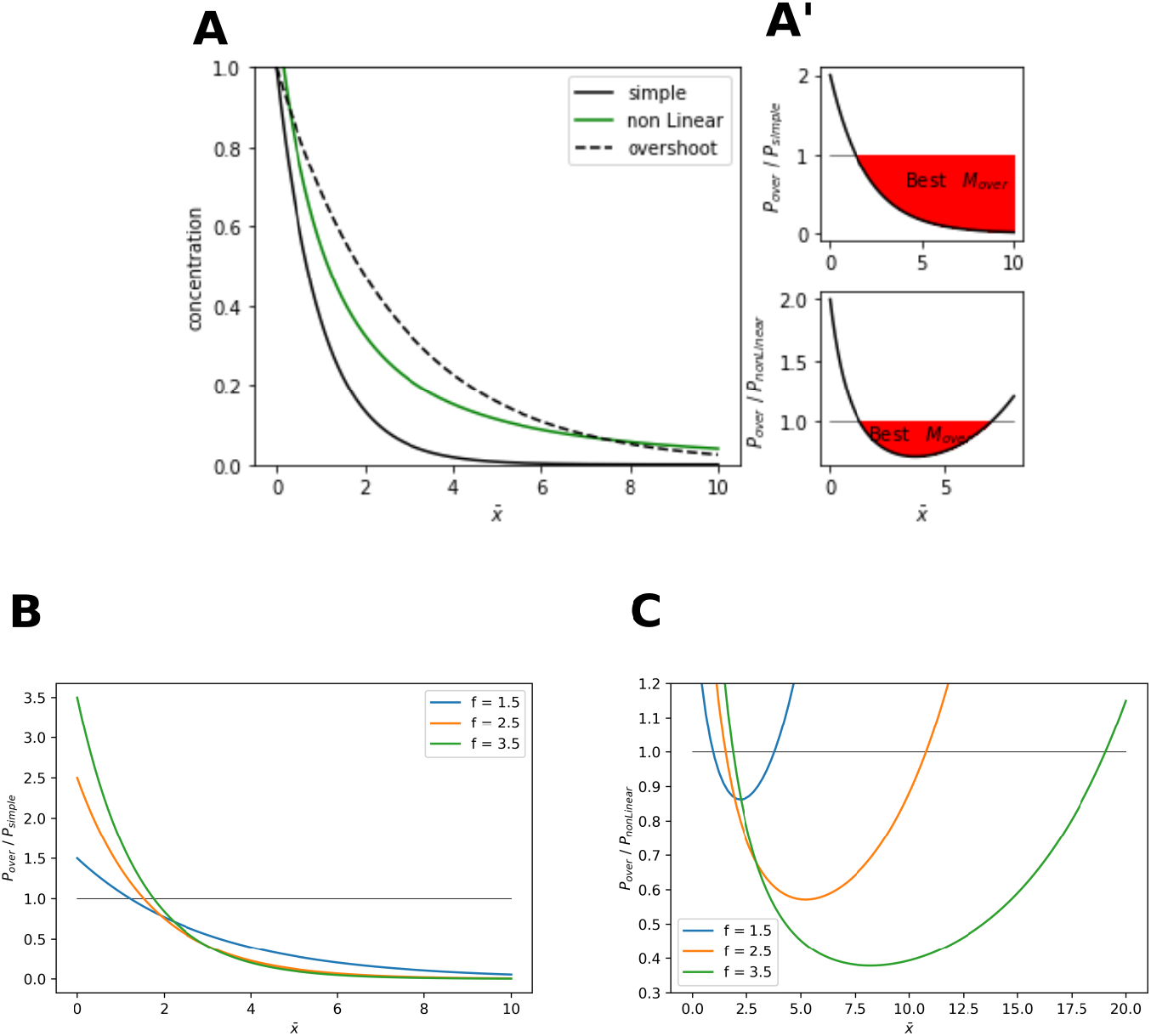
(Supplementary figure to Figure 2). Comparison of precision coefficients for steady-state linear and non-linear decay gradients and the overshoot profile. A. Gradients resulting from a simple diffusion/linear degradation model at the steady-state (black), non-linear degradation model at steady-state (*F* (*M*) =^2^; green; see equation (10) and section 2.3 in the Supplementary Information). A’. Pair-wise comparison of the precision coefficients for the models in A. Red areas indicate the positions where the ratios below 1. Note that there is a region where the overshoot model is more precise than the non-linear decay model. B, C. changes in the ratios computed in A’ when the overshoot is approximated by an exponential function given by *λ*_*over*_ = *f × λ*_*SS*_ for three different values of *f*. Note that as *f* increases, the territory where the overshoot model is more precise increases. 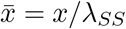.

**Supplementary figure 2:**
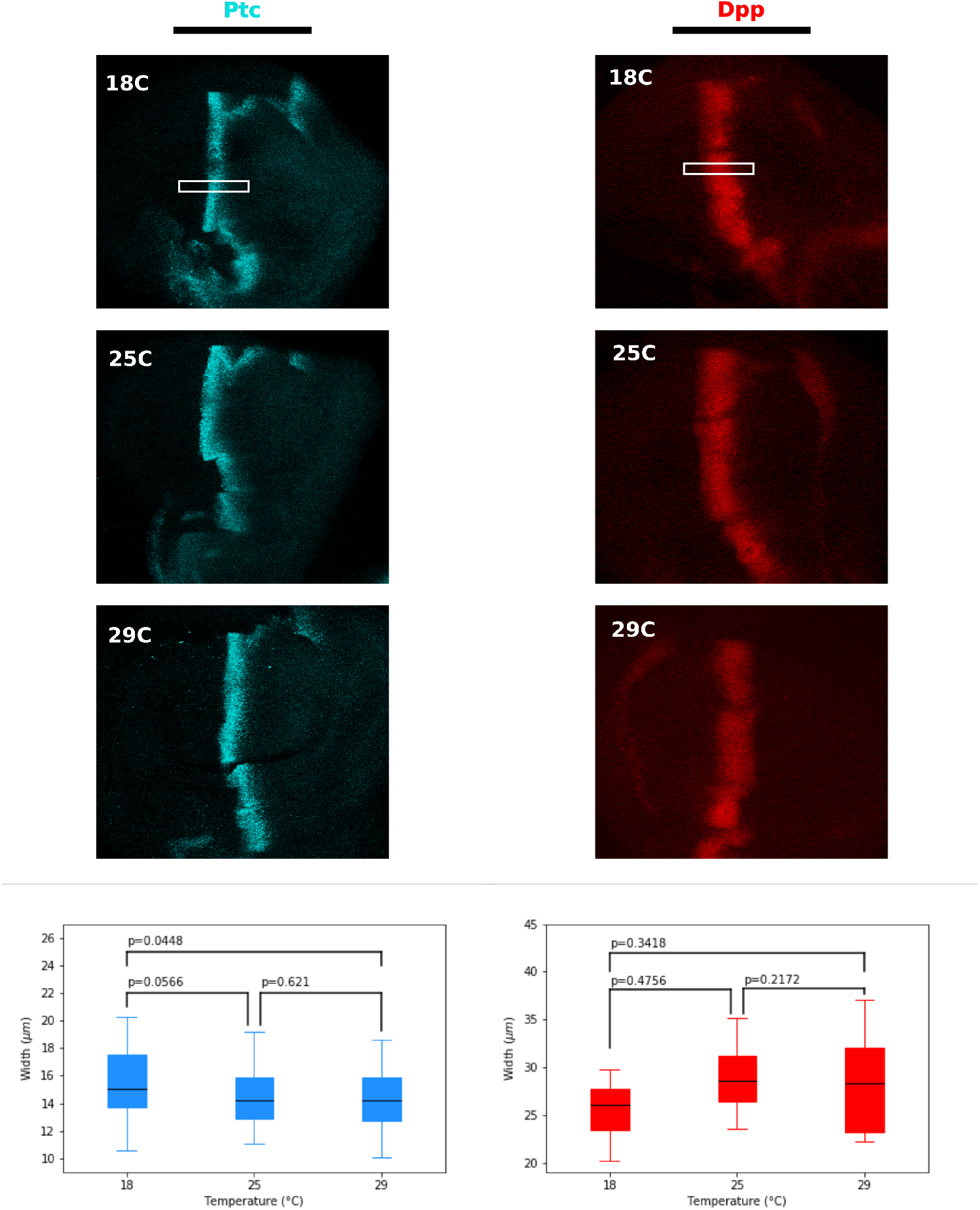
(Supplementary figure to Figure 6). Patterns of Hh target genes are not significantly affected by temperature. First column, Ptc; second column Dpp-GFP. Below, measures of width pattern in marked rectangle. The graphs underneath compare the widths of the corresponding patterns at different temperatures. p-values are computed after a Student t-test (when the data had normal distributions) or Mann-Whitney U (when the data did not have normal distributions) of each pair of datasets.

