## Supplementary information for "Dynamic readout of the Hh gradient in the Drosophila wing disc reveals pattern-specific tradeoffs between robustness and precision"

### 1 Proof that all thresholds shift uniformly at the steady-state

In this section, we show that at the steady-state, different morphogen diffusion models have the same effect on their target genes upon perturbations in morphogen production rates.

#### 1.1 Simple diffusion model

First, we analyze the model of simple diffusion and linear degradation at the steady-state:

$$\frac{d^2M}{dx^2} - \lambda^2 M = 0, \tag{1}$$

where $M$ is the concentration of the morphogen and $\lambda^2$ is the ratio between the diffusion coefficient ($D$) and the degradation ($\beta$). The model is subject to the following boundary conditions:

$$\begin{aligned} &B.C.1. \quad M(0) = M_0, \\ &B.C.2. \quad \lim_{x\to\infty} M(x) = 0. \end{aligned} \tag{2}$$

The analytical solution of this problem is:

$$M(x) = M_0 e^{-\lambda x}. \tag{3}$$

where $M_0$ is the ligand concentration at the source position $x = 0$. Now consider the perturbation, $M_0 \to \tilde{M}_0$, then perturbed solution will be:

$$\begin{aligned} \tilde{M}(x) &= \tilde{M}_0 \exp(-\lambda x) \\ &= M_0 \exp\left(\ln\left(\frac{\tilde{M}_0}{M_0}\right)\right) \exp(-\lambda x) \\ &= M_0 \exp\left(-\lambda\left(x - \frac{1}{\lambda}\ln\left(\frac{\tilde{M}_0}{M_0}\right)\right)\right) \\ &= M(x-a), \quad a = \frac{1}{\lambda}\ln\left(\frac{\tilde{M}_0}{M_0}\right). \end{aligned} \tag{4}$$

Therefore, all the territories defined by thresholds of this gradient will be displaced by the same amount $a$.

#### 1.2 Diffusion model with a general degradation function

Consider the following model originally introduced by Eldar et al. (2003):

$$\frac{d^2M}{dx^2} - \lambda^2 F(M) = 0, \tag{5}$$

subject to the boundary condition:

$$M(0) = M_b^{(1)}, \tag{6}$$

whose solution is:

$$x(L; M_b) = \lambda \int_M^{M_b^{(1)}} \frac{dU}{\sqrt{E(U)}}; \quad E(M) = 2\int_0^M F(U)dU. \tag{7}$$

Again, the source is modeled as a boundary condition, so let us consider a perturbation $M_b^{(1)} \to M_b^{(2)}$. The perturbed solution is:

$$\begin{aligned}\tilde{x}(M; M_b^{(2)}) &= \lambda \int_M^{M_b^{(2)}} \frac{dU}{\sqrt{E(U)}} \\ &= \lambda \int_M^{M_b^{(1)}} \frac{dU}{\sqrt{E(U)}} + a \\ &= x(M; M_b^{(1)}) + a,\end{aligned} \tag{8}$$

where $a$ is given by $\lambda \int_{M_b^{(1)}}^{M_b^{(2)}} \frac{dU}{\sqrt{E(U)}}$. Then

$$\tilde{M}(\tilde{x}(M; M_b^{(2)})) = M(x(M; M_b^{(1)}) + a) \tag{9}$$

a uniform displacement by the amount $a$ of the entire profile.

#### 1.3 Diffusion model with a source explicitly in the equation

So far we have analyzed models where the source is represented as a boundary condition. This raises the question of what happens in the case where the source appears explicitly in the equations. Consider the model:

$$D\frac{d^2M}{dx^2} + \theta^-(x)\alpha - \beta M = 0 \tag{10}$$

where $\theta^-(x) = 1$ for $x < 0$ and 0 otherwise, this is, we have the case of a source that produces ligand at a uniform rate in the negative region of the $x$-axis.

We consider the following boundary conditions (B.C.),

$$\begin{aligned} lim_{x\to-\infty} M = 0, \\ lim_{x\to\infty} M = 0. \end{aligned} \tag{11}$$

And let us assume continuity of $M$ and its derivative at $x = 0$:

$$\begin{aligned} M^+(0) &= M^-(0) \\ \left.\frac{dM^+}{dx}\right|_{x=0} &= \left.\frac{dM^-}{dx}\right|_{x=0}, \end{aligned} \tag{12}$$

where $M^-$ is the solution for $x < 0$ and $M^+$ for $x > 0$. The solution of the problem given by equations (10)-(12) is:

$$M(x) = \begin{cases} -\frac{\alpha}{2\beta} e^{\lambda x} + \frac{\alpha}{\beta} & x \leq 0, \\ \frac{\alpha}{2\beta} e^{-\lambda x} & x > 0. \end{cases} \tag{13}$$

Note that $M(x=0) = \frac{\alpha}{2\beta}$ and that for $x > 0$, the solution is the same as in the case where the source is modeled as a boundary condition (eq. (1)). Therefore, we conclude that:

- Even when the source is modeled explicitly, the ligand profile only depends on the concentration of the ligand at the border of the source.
- All the territories defined by the steady-state profile are displaced by the same amount.

#### 1.4 Simple model of the Hedgehog (Hh) morphogen

Consider the following model of the Hh morphogen in the *Drosophila* wing disc:

$$\frac{\partial Hh}{\partial t} = D\frac{\partial^2 Hh}{\partial x^2} + \theta(x)\alpha_{Hh} - \gamma_{Hh\_Ptc}(Hh) \cdot (Ptc) - \beta_{Hh}Hh, \tag{14}$$

where $\theta(x)\alpha_{Hh}$ represents the source of Hh in the posterior compartment of the wing disc and the term $\gamma_{Hh\_Ptc}Hh \times Ptc$ represents ligand binding with the Patched (Ptc) receptor. We can consider the following term as a generalized degradation function:

$$\beta(x,t) = (\gamma_{Hh\_Ptc}Ptc - \beta_{Hh})\, Hh. \tag{15}$$

Then, at steady-state $\beta_{steady-state}(x) = \gamma_{Hh\_Ptc}Ptc_{steady-state}(x) - \beta_{Hh}$ . At steady-state we expect that $Ptc_{steady-state}$ forms a uniform expression pattern over a stripe of anterior cells abutting the Anterior-Posterior (AP) border. Therefore, at steady-state $Ptc_{steady-state}$ can be modeled as a step function, and the binding of Hh to Ptc as a linear degradation given by:

$$D\frac{d^2 Hh}{dx^2} + \theta(x)^{-}\alpha - \beta_{steady-state}Hh = 0, \tag{16}$$

where,

$$\beta_{steady-state}(x) = \begin{cases} \beta^*, & 0 < x < b \\ \beta, & \text{otherwise} \end{cases} \tag{17}$$

where $b$ is the width of Ptc stripe, $\theta(x)$ is a step function centered at zero and $\beta^* > \beta$.

This equation can be solved in three territories: the first territory is $x > 0$, and the solution there will be denoted by $Hh_{anterior}$. The second territory is $0 < x < b$ and solution will be denoted by $Hh_{stripe}$. The third territory is $x > b$, and the solution will be denoted by $Hh_{beyondPtc}$.

For $x < 0$,

$$Hh_{\text{anterior}}(x) = P + e^{\lambda_1 x} + \alpha/\beta, \tag{18}$$

where $\lambda_1^2 = \beta/D$ and $P$ is a constant that will be determined by boundary conditions. For $0 < x < b$,

$$Hh_{\text{stripe}}(x) = Ae^{\lambda_2 x} + Be^{-\lambda_2 x}, \tag{19}$$

where $\lambda_2^2 = \beta^*/D$ and $A$, $B$ are constants that will be determined by boundary conditions. For $x \geq b$,

$$Hh_{beyondPtc}(x) = Ce^{-\lambda_1 x}, \tag{20}$$

where $C$ will be determined by boundary conditions.

By imposing continuity of functions at $x = 0$ and $x = b$ we obtain,

$$P = \frac{\frac{\alpha}{\beta}\left[e^{-\lambda^- b}\left(\frac{\lambda_2}{\lambda_1} - 1\right) - e^{\lambda^+ b}\left(1 + \frac{\lambda_2}{\lambda_1}\right)\right]}{e^{\lambda^+ b}\left(1 + \frac{\lambda_1}{\lambda_2}\right)\left(1 + \frac{\lambda_2}{\lambda_1}\right) - e^{-\lambda^- b}\left(1 - \frac{\lambda_1}{\lambda_2}\right)\left(\frac{\lambda_2}{\lambda_1} - 1\right)}, \tag{21}$$

where

$$\begin{aligned} \lambda^+ &= \lambda_2 + \lambda_1, \\ \lambda^- &= \lambda_2 - \lambda_1. \end{aligned} \tag{22}$$

Then,

$$A = \frac{P}{2}\left(1 + \frac{\lambda_1}{\lambda_2}\right) + \frac{\alpha}{2\beta}, \tag{23}$$

$$B = \frac{P}{2}\left(1 - \frac{\lambda_1}{\lambda_2} + \frac{\alpha}{2\beta}\right), \tag{24}$$

$$C = Ae^{\lambda^+ b} + Be^{-\lambda^- b}. \tag{25}$$

Therefore, for $x > 0$,

$$Hh(x) = \begin{cases} Hh_{\text{stripe}} = Ae^{\lambda_2 x} + Be^{-\lambda_2 x}, & 0 < x < b \\ Hh_{beyondPtc} = Ce^{-\lambda_1 x}, & x \geq b. \end{cases} \tag{26}$$

Note that upon a perturbation $\alpha \to \tilde{\alpha}$, $Hh_{\text{stripe}} \to \tilde{H}h_{\text{stripe}}$, where

$$\tilde{H}h_{\text{stripe}} = Ae^{\lambda_2\left[x + \frac{1}{\lambda_2}\ln\left(\frac{\tilde{\alpha}}{\alpha}\right)\right]} + Be^{-\lambda_2\left[x - \frac{1}{\lambda_2}\ln\left(\frac{\tilde{\alpha}}{\alpha}\right)\right]}, \tag{27}$$

and $Hh_{beyondPtc} \to \tilde{H}h_{beyondPtc}$ , where,

$$\tilde{H}h_{beyondPtc} = Ce^{-\lambda_1\left[x - \frac{1}{\lambda_1}\ln\left(\frac{\tilde{\alpha}}{\alpha}\right)\right]}. \tag{28}$$

We conclude that all territories defined by $Hh_{beyondPtc}$ are shifted by the same amount, $\frac{1}{\lambda_1}\ln\left(\frac{\tilde{\alpha}}{\alpha}\right)$, upon variations in $\alpha$.

### 2 Dynamic interpretation of morphogen profile implies differential robustness

#### 2.1 Simple diffusion model

Consider the time-independent model given by,

$$\frac{\partial M}{\partial t} = D\frac{\partial^2 M}{\partial x^2} - \beta M, \tag{29}$$

subject to the following boundary and initial conditions:

$$\begin{aligned} \text{B.C.1.} \quad & M(0,t) = M_0, \\ \text{B.C.2.} \quad & \left.\frac{\partial M}{\partial x}\right|_{x=l} = 0, \\ \text{I. C.} \quad & M(x,0) = M_0 F(x), \end{aligned} \tag{30}$$

where $l$ defines the length of the tissue. To solve this equation we split the solution in two parts:

$$M(x,t) = M_{ts}(x) + M_{ss}(x,t), \tag{31}$$

where $M_{ts}$ means transient solution and $M_{ss}$ steady-state solution, respectively. We known that $M_{ss}$ satisfy

$$\begin{aligned} D\frac{\partial^2 M_{ss}}{\partial x^2} - \beta M_{ss} &= 0, \\ \text{B.C.1.} \quad M_{ss}(0,t) &= M_0, \\ \text{B.C.2.} \quad \left.\frac{\partial M_{ss}}{\partial x}\right|_{x=l} &= 0. \end{aligned} \tag{32}$$

Replacing the $M$ split sum we have

$$\begin{aligned} \frac{\partial M_{ts}}{\partial t} &= D\frac{\partial^2 M_{ts}}{\partial x^2} - \beta M_{ts}, \\ \text{B.C.1.} \quad & M_{ts}(0,t) = 0, \\ \text{B.C.2.} \quad & \left.\frac{\partial M_{ts}}{\partial x}\right|_{x=l} = 0, \\ \text{I. C.} \quad & M(x,0) = \begin{cases} M_0 & \text{if } 0 \le x \le a, \\ 0 & x > a \end{cases} \end{aligned} \tag{33}$$

**Steady-state solution.** The general solution for $M_{ss}$ equation is given by:

$$M_{ss}(x) = Ae^{x/\lambda} + B^{-x/\lambda}; \quad \lambda = \sqrt{\frac{D}{\beta}}, \tag{34}$$

where $A$ and $B$ are determined by the B. C. Applying the B.C.s we get:

$$M_{ss}(x) = M_0 \left[ \frac{e^{-l/\lambda}}{\cosh(l/\lambda)} \sinh(x/\lambda) + e^{-x/\lambda} \right]. \tag{35}$$

**Transient solution.** Now we solve the equation:

$$\begin{aligned} \frac{\partial M_{ts}}{\partial t} &= D \frac{\partial^2 M_{ts}}{\partial x^2} - \beta M_{ts}, \\ \text{B.C.1.} \quad & M_{ts}(0, t) = 0, \\ \text{B.C.2.} \quad & \left. \frac{\partial M_{ts}}{\partial x} \right|_{x=l} = 0, \\ \text{I. C.} \quad M(x, 0) &= \begin{cases} M_0 & \text{if } 0 \le x \le a, \\ 0 & x > a. \end{cases} \end{aligned} \tag{36}$$

We known that the equation can be solved by separation of variables because it is a homogeneous equation with homogeneous B.C.s. Consider the solution $M_{ts}(x, t) = M_x(x) \cdot M_t(t)$, so

$$\begin{aligned} \frac{dM_t}{dt} + k^2 D \cdot M_t = 0, & \quad \text{temporal equation.} \\ \frac{d^2 M_x}{dx^2} - (\lambda^2 - k^2) M_x = 0, & \quad \text{spatial equation,} \end{aligned} \tag{37}$$

where $k$ is a constant of separation of variable that will be determined below. The temporal equation have the solution

$$M_t(t) = e^{-k^2 Dt}. \tag{38}$$

The spatial equation have the general solution,

$$\frac{d^2 M_x}{dx^2} = A e^{\sqrt{1/\lambda^2 - k^2} x} + B e^{-\sqrt{1/\lambda^2 - k^2} x}, \tag{39}$$

but we have two cases, $1/\lambda^2 - k^2 > 0$, and the second $1/\lambda^2 - k^2 < 0$. The first case leads to a contradiction, so we focus on the last.

Let $\Lambda^2 = k^2 - 1/\lambda^2$, so $\sqrt{1/\lambda^2 - k^2} = i\Lambda$

$$M_x(x) = A e^{i\Lambda x} + B e^{-i\Lambda x}. \tag{40}$$

We select a real solution:

$$M_x(x) = A \sin(\Lambda x) + B \cos(\Lambda x). \tag{41}$$

Then,

$$M(x, t) = e^{-k^2 Dt} \left[ A \sin(\Lambda x) + B \cos(\Lambda x) \right]. \tag{42}$$

The second B.C. leads to

$$M(x,t) = Ae^{-k^2 Dt}\left[\sin(\Lambda x) - \frac{\cos(l\Lambda)}{\sin(l\Lambda)}\cos(\Lambda x)\right]. \quad (43)$$

The firts B.C. leads to $\cos(\Lambda M) = 0$, or $\Lambda M = \frac{\pi}{2} + n\pi, \quad n = 0, 1, 2, \ldots,$

$$\begin{aligned} M_n(x,t) &= A_n e^{-k_n^2 Dt}\sin(\Lambda_n x), \\ \Lambda_n &= \frac{\pi + 2n\pi}{2l}, \\ k_n^2 &= \left(\frac{\pi + 2n\pi}{2l}\right)^2 + 1/\lambda^2. \end{aligned} \quad (44)$$

We propose a particular solution of the form

$$M_{ts}(x,t) = \sum_{n=0}^{\infty} A_n e^{-k_n^2 Dt}\sin(\Lambda_n x). \quad (45)$$

The I. C. leads to

$$\sum_{n=0}^{\infty} A_n \sin(\Lambda_n x) = M_0 F(x), \quad (46)$$

using the orthogonality relations

$$\frac{2}{l}\int_0^l \sin(\Lambda_n x)\sin(\Lambda_m x)dx = \begin{cases} 1 & \text{if } n = m, \\ 0 & \text{otherwise.} \end{cases} \quad (47)$$

we obtain

$$A_n = \frac{2M_0}{l}\int_{-100}^{0} F(x)\sin(\Lambda_n x)dx. \quad (48)$$

Therefore,

$$M(x,t) = M_0\left[\frac{e^{-l/\lambda}}{\cosh(l/\lambda)}\sinh(x/\lambda) + e^{-x/\lambda} + \sum_{n=0}^{\infty}\hat{A}_n e^{-k_n^2 Dt}\sin(\Lambda_n x)\right], \quad (49)$$

where $\hat{A}_n = A_n/M_0$. Note that $M(x,t)$ only depend on $M_0$ by a multiplication.

**Fourier series of $M_{SS}$.** Since the set of functions $\{\sin(\Lambda_n x) | n = 0, 1, 2, \ldots\}$ form a base in the same space of the steady-state solution we can compute the Fourier series of $M_{SS}$.

Let,

$$M_{SS} = \sum_n B_n \sin(\Lambda_n x), \quad (50)$$

then

$$B_n = \frac{2}{l}\frac{M_0\Lambda_n}{\Lambda_n^2 + 1}. \quad (51)$$

##### 2.1.1 Coefficient of precision in a morphogen model

We define *coefficient of precision* of the morphogen gradient by:

$$\mathcal{P} = \left| \frac{dx}{dM} \right| = \left| \frac{1}{\frac{dM(x,t1)}{dx}} \right| \tag{52}$$

In order to known the $\mathcal{P}$ of our ligand function, $M(x,t)$ from eq. (29), we need to determine the slope of the tangent line to $M(x,t_1)$ for a given time $t_1$. Since

$$\left.\frac{\partial M}{\partial x}\right|_{t_1} = \frac{dM_{SS}}{dx} + \left.\frac{\partial M_{ts}}{\partial x}\right|_{t_1}, \tag{53}$$

we compare $\mathcal{P}$ of transient and steady-state solutions by knowing the relation ($>, = or <$) between $\left|\frac{\partial M_{tr}}{\partial x} + \frac{dM_{SS}}{dx}\right|$ and $\left|\frac{dM_{SS}}{dx}\right|$. They cannot be equal because $\lim_{t\to\infty} \frac{\partial M_{tr}}{\partial x} = 0$ in such away that

$$\lim_{t\to\infty} \frac{\partial M(x,t)}{\partial x} = \frac{dM_{SS}(x)}{dx}. \tag{54}$$

This last result implies that $\mathcal{P}$ of transient solution tend to the steady-state $\mathcal{P}$.

By the other hand, if $\frac{\partial M_{tr}}{\partial x}$ and $\frac{dM_{SS}}{dx}$ have the same sign then $\left|\frac{\partial M_{tr}}{\partial x} + \frac{dM_{SS}}{dx}\right| > \left|\frac{dM_{SS}}{dx}\right|$. When one differentiate the two functions one can see that both have the same sign. Then, for the model defined by eq. (29), and for each time, the transient solution is more precise than the steady-state solution.

#### 2.2 Simplified dynamic model of Hh signaling

Consider the simplified dynamic model of a Hh gradient, given by the equation:

$$\begin{aligned} \frac{\partial Hh}{\partial t} &= D\frac{\partial^2 Hh}{\partial x^2} - \beta(t)Hh, \\ \beta(t) &= \begin{cases} \beta_{early}, & 0 \le t \le t_A \\ \beta_{late}, & t > t_A \end{cases} \qquad \beta_{early} < \beta_{late} \end{aligned} \tag{55}$$

subject to the boundary condition,

$$\text{B. C.} \quad Hh(x=0,t) = Hh_0 \tag{56}$$

where $t_A$ is the time at which the rate of degradation change between initial rate $\beta_{ealy}$ and final rate $\beta_{late}$. This model attempts to account for the upregulation of *ptc* by Hh. $\beta_{early}$ represents the basal degradation of Hh. When the ligand induces the expression of its receptor, it results in an increased degradation of the morphogen, $\beta_{late}$. If we consider the generalized degradation function, eq. (15), we can write $\beta_{late}$ as $\gamma_{Hh\_Ptc}Ptc_{steady-state} - \beta_{Hh}$. The transitory states of this model will move between the pseudo-state corresponding to $\beta_{early}$,

$$Hh_{pseudoSS} = Hh_0 e^{-x/\lambda_{early}}, \tag{57}$$

with $\lambda^2_{early} = D/\beta_{early}$ and the real steady-state

$$Hh_{realSS} = Hh_0 e^{-x/\lambda_{late}}, \tag{58}$$

with $\lambda^2_{late} = D/\beta_{late}$. Since

$$\lambda_{early} > \lambda_{late}, \tag{59}$$

the reach of $Hh_{pseudoSS}$ is greater than the reach of $Hh_{realSS}$. We define the *overshoot* gradient (in the context of Nahmad and Stathopoulos, 2009) as the transient state with the longest reach.

Note that the solution (49) can be applied to our model of Hh (eq. (55)) for $t < t_A$ and $t > t_A$ independently.

For $t < t_A$ we have:

$$Hh_{early}(x,t) = Hh_0 \left[ \frac{e^{-l/\lambda_{early}}}{\cosh(l/\lambda_{early})} \sinh(x/\lambda_{early}) + e^{-x/\lambda_{early}} + \sum_{n=0}^{\infty} \hat{A}_{early\_n} e^{-k_n^2 Dt} \sin(\Lambda_{early\_n} x) \right]. \tag{60}$$

Then, the overshoot is given by $Hh_{early}(x, t_A)$. Taken the overshoot as initial condition for late times we have (for $t > t_A$),

$$Hh_{late}(x,t) = Hh_0 \left[ \frac{e^{-l/\lambda_{late}}}{\cosh(l/\lambda_{late})} \sinh(x/\lambda_{late}) + e^{-x/\lambda_{late}} + \sum_{n=0}^{\infty} \hat{A}_{late\_n} e^{-k_n^2 Dt} \sin(\Lambda_{late\_n} x) \right]. \tag{61}$$

Again, we note that $Hh(x,t)$ only appears multiplied by the source concentration, $Hh_0$.

##### 2.2.1 Coefficient of precision of dynamical model of Hh signaling

Let us calculate $\mathcal{P}$ for the pseudoSS and for realSS. Replacing eqs. (60) and (61) in to definition (eq. (52)):

$$\begin{aligned} \mathcal{P}_{pseudoSS} &= \frac{\lambda_{early}}{T}, \\ \mathcal{P}_{realSS} &= \frac{\lambda_{late}}{T}, \end{aligned} \tag{62}$$

where $T$ is the threshold at which territories are defined. From eq. (59) we have that $\mathcal{P}_{realSS} < \mathcal{P}_{pseudoSS}$. Likewise, from eq.(54) we have that $\mathcal{P}_{overshoot} < \mathcal{P}_{pseudoSS}$. Finally, since we are considering $t_A$ in such away that the reach of the overshoot is bigger than the reach of realSS, and due to after the overshoot transient solutions tend to realSS, we have that $\frac{\partial Hh_{overshoot}}{\partial x} > \frac{dHh_{realSS}}{dx}$, Therefore,

$$\mathcal{P}_{realSS} < \mathcal{P}_{overshoot} < \mathcal{P}_{psuedoSS}. \tag{63}$$

##### 2.2.2 Robustness of the simplified model of Hh signaling

We define the coefficient of robustness of the territories defined by the morphogen gradient with respect to the source $Hh_0$ through

$$\mathcal{R} \equiv \left| \frac{dx}{dHh_0} \right| = \mathcal{P} \times \left| \frac{dHh}{dHh_0} \right| \tag{64}$$

Here, since $H(x,t)$ only appears multiplied by $Hh_0$ (see equations (60) and (61)) becomes relevant. So if we define $Hh(x,t) = Hh_0 \times \hat{H}h(x,t)$, then $\frac{dHh}{dHh_0} = \hat{H}h$. On the other hand, the dynamic model defines different territories at the same concentration, so $\hat{H}(x,t) = \frac{T}{Hh_0}$ , where $T$ is the concentration at which the territories are defined. Then, for the simplified model:

$$\mathcal{R} = \frac{Hh_0}{T} \times \mathcal{P} \tag{65}$$

Equation (65) establishes a direct relationship between $\mathcal{R}$ and $\mathcal{P}$. Therefore from eq. (63):

$$\mathcal{R}_{realSS} < \mathcal{R}_{overshoot}, \tag{66}$$

*i.e.*, the overshoot has lower robustness than the steady-state.

#### 2.3 Coefficients of precision for different models

Now we are going to compare $\mathcal{P}$ for our dynamical and steady-state models. Let $M_{simple}$, $M_{nonLinear}$ and $M_{over}$ the solutions for the models (3), non-linear with $F(M) = M^2$ (see Eldar et al. (2003)) and our dynamical model (55), respectively. For $t_A$ large enough we can approximate the overshoot by a pseudoSS solution $M_{over} \approx M_0 e^{-x/\lambda_{over}}$. We define the *$\lambda$-increment factor*, $f$, in such away

$$\lambda_{over} = f \times \lambda_{realSS}, \tag{67}$$

then

$$\frac{\mathcal{P}_{over}}{\mathcal{P}_{simple}} = f e^{(1/f-1)\bar{x}}, \tag{68}$$

$$\frac{\mathcal{P}_{over}}{\mathcal{P}_{nonLinear}} = \frac{2f}{\sqrt{6}M_0} \frac{e^{\bar{x}/f}}{\left( \frac{\bar{x}}{\sqrt{6}} + \left( \frac{2}{\sqrt{6}M_0} \right)^{1/3} \right)^3}, \tag{69}$$

where $\bar{x} = x/\lambda_{realSS}$. Comparison of equations (68) and (69) for different values of $f$ is showed in Supplementary figure 1A'. In the comparison with the simple model we can observe two territories: one close to the source, where the steady-state model is more precise ($\mathcal{P}_{over}/\mathcal{P}_{simple} > 1$), and another from which the overshoot model is more precise ($\mathcal{P}_{over}/\mathcal{P}_{simple} < 1$, Supplementary figure 1A' above). In the comparison with the nonlinear model, we can observe three territories: one close to the source, where the nonlinear model is more precise than the overshoot, another intermediate one where the overshoot model is more precise (which corresponds to the location where *dpp* is positioned in our system), and a last one where again the nonlinear model is more precise (see Supplementary figure 1A' below). In the case of Hh signaling in the *Drosophila* wing disc, this last territory is outside the range of Hh signaling (Kicheva and González-Gaitán, 2008). Furthermore the territory where the overshoot model is more precise grows as $f$ increases (see Supplementary figure 1A-C).

### 3 Hh signaling model

A more explicit model of Hh signaling is decribed in Box 3. At the steady-state, the equations can be reduced to a single equation in each compartment (Nahmad and Stathopoulos, 2009):

$$D\frac{d^2Hh_{SS}}{dx^2} + S^+(x)\alpha_{Hh} - \frac{\chi S^-(x)Hh_{SS}}{\gamma_{Hh\_Ptc}Hh + \beta_{Ptc}}\left[\alpha_{ptc0} + \frac{\alpha_{ptc}Hh_{SS}^{nm}}{\eta^m\left(k^n + Hh_{SS}^n\right)^m + S^-(x)Hh_{SS}^{nm}}\right] - \beta_{Hh}Hh_{SS} = 0, \tag{70}$$

where,

$$k = \frac{k_{Signal}\beta_{Hh\_Ptc}}{\gamma_{Hh\_Ptc}}, \tag{71}$$

$$\eta = \frac{k_{ptc}\beta_{Signal}}{\alpha_{Signal}}, \tag{72}$$

$$\chi = \frac{T_{Ptc}\gamma_{Hh\_Ptc}}{\beta_{ptc}}. \tag{73}$$

We use the subscript SS to denote steady-state concentrations, but when it does not cause confusion we ignore the subscript. The steady-state equation satisfies the boundary condition,

$$\left.\frac{dHh_{SS}}{dx}\right|_{x=-100} = \left.\frac{dHh_{SS}}{dx}\right|_{x=100}. \tag{74}$$

#### 3.1 Signal depends only on Hh at the steady-state

At the steady-state, simple sustitution results in:

$$Signal(x) = \begin{cases} 0, & x < 0, \\ \frac{\alpha_{Signal}}{\beta_{Signal}}\frac{Hh^n(x)}{k^n + Hh^n(x)}, & x \geq 0, \end{cases} \tag{75}$$

where $k = \frac{k_{Signal}\beta_{Hh\_Ptc}}{\gamma_{Hh\_Ptc}}$.

Notice that at Steady-State, equation for $Hh$ is reduced to

$$D\frac{d^2Hh}{dx^2} + S^+(x)\alpha_{Hh} - \beta_{SS}(x)Hh = 0, \tag{76}$$

where $\beta_{SS}(x) = \gamma_{Hh\_Ptc} Ptc_{SS} + \beta_{Hh}$, the same equation that (16). In section 1.2 we proved which under perturbations on $\alpha$ all the territories are shifted by the same amount. since $Signal$ only depends on $Hh$ at the steady-state, for $x \geq 0$, a shift of $Hh$ by the amount $a$ means a shift by the same amount in $Signal$,

$$Signal(x-a) = \frac{\alpha_{Signal}}{\beta_{Signal}} \frac{Hh^n(x-a)}{k^n + Hh^n(x-a)}. \tag{77}$$
